# How to select the best zero-shot model for the viral proteins?

**DOI:** 10.1101/2024.10.06.616860

**Authors:** Yuanxi Yu, Fan Jiang, Bozitao Zhong, Liang Hong, Mingchen Li

## Abstract

Predicting the fitness of viral proteins holds notable implications for understanding viral evolution, advancing fundamental biological research, and informing drug discovery. However, the considerable variability and evolution of viral proteins make predicting mutant fitness a major challenge. This study introduces the ProPEC, a Perplexity-based Ensemble Model, aimed at improving the performance of zero-shot predictions for protein fitness across diverse viral datasets. We selected five representative pretrained language models (PLMs) as base models. ProPEC, which integrates perplexity-weighted scores from these PLMs with GEMME, demonstrates superior performance compared to individual models. Through parameter sensitivity analysis, we highlight the robustness of perplexity-based model selection in ProPEC. Additionally, a case study on T7 RNA polymerase activity dataset underscores ProPEC’s predictive capabilities. These findings suggest that ProPEC offers an effective approach for advancing viral protein fitness prediction, providing valuable insights for virology research and therapeutic development.

**TOC Graphic:** 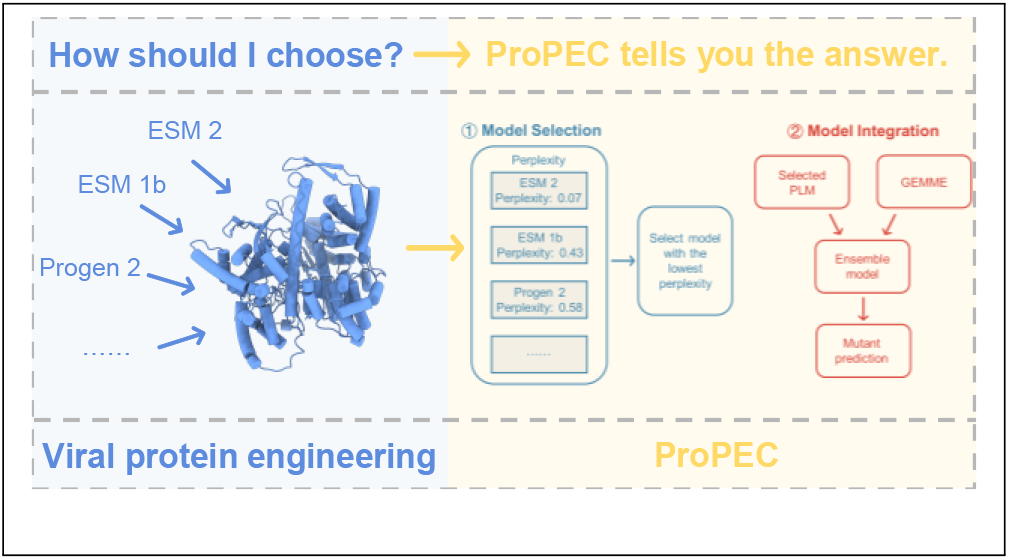

## Introduction

Protein mutation prediction is pivotal in deciphering the consequences of genetic variations on protein function,^1,2^ stability,^3,4^ and interactions.^5,6^ This understanding is crucial for applications such as drug design,^7^ disease mechanism elucidation,^8^ and therapeutic intervention development.^9,10^ While traditional methods like GEMME^11^ and EVE,^12^ which rely on evolutionary conservation and multiple sequence alignment (MSA) data, have been effectively used for these predictions. In recent years, deep learning models have significantly advanced this field by leveraging large-scale protein sequence data to predict mutation effects.^13^ For example, pretrained language models (PLMs) like ESM-2, ^14^ Progen2,^15^ and SaProt^16^ have shown remarkable accuracy on zero-shot predictions of protein mutant fitness in absence of experimental data. Since these models fundamentally represent statistical characteristics of natural protein sequences found in nature, their zero-shot likelihood scores for mutation fitness essentially measure how similar a mutant protein sequence is to natural proteins or a particular protein family.^17,18^

In the context of natural proteins, viral proteins stand out as a particularly unique category.^19^ They exhibit distinctive properties such as lower contact density, a higher proportion of disordered and coil regions, and greater structural flexibility.^20,21^ These characteristics enable viral proteins to interact effectively with host components and adapt swiftly to environmental changes. Unlike proteins from thermophilic or mesophilic organisms, which rely on high stability (threshold robustness) to buffer against mutations, viral proteins employ a strategy of loose packing and partial disorder. ^19^ The fitness of mutants in viral proteins is often more subtle and harder to capture compared to other proteins, reflecting their high mutation rates and rapid evolutionary dynamics. ^19,22^ Therefore, predicting the fitness of viral protein mutants in the absence of sufficient experimental data and domain knowledge has become an attractive challenge for zero-shot approaches.

For zero-shot predictions on viral proteins, although some PLMs have achieved promising results, their performance tends to vary significantly across different proteins, making them insufficiently robust.^23^ For example, according to the investigation on Proteingym,^24^ a benchmark including an extensive deep mutational scanning (DMS) datasets, the PLMs can achieve an average score of 0.485 on non-viral protein datasets. However, on viral datasets,the best PLM only reaches a score of 0.432, which is lower than the best performer: GEMME with the score of 0.469. As a result, ensemble methods are often employed in practical applications.^25–27^ However, current ensemble approaches are typically suboptimal, as they directly combine or stack features or contributions from different models. This method overlooks the varying biases each model may have toward the wild-type protein, failing to account for the strengths and weaknesses of individual models. To overcome this limitation, we present ProPEC (Protein Perplexity Ensemble Classifier), an ensemble method that combines perplexity-based predictions from multiple PLMs with GEMME. By leveraging perplexity—a measure of a model’s uncertainty in predicting a sequence^28^ we weight the contributions of different PLMs based on their performance on wild-type proteins. Our findings reveal a negative correlation between a model’s perplexity on wild-type sequences and its fitness prediction accuracy for protein mutants, indicating that models with lower perplexity (higher confidence) tend to make more accurate predictions. Importantly, calculating perplexity does not require any experimental cost, allowing us to effectively select the best-performing models without domain knowledge or additional resources. The predictions from perplexity-selected PLMs are then combined with GEMME, leveraging the statistical characteristics from deep learning models alongside the evolutionary information encapsulated in MSA-based methods.

In this study, we introduce ProPEC, an ensemble method that combines perplexity-weighted predictions from multiple pretrained language models with GEMME, for selecting the best zero-shot models for viral protein mutation prediction. We systematically evaluate ProPEC’s performance across 23 viral protein datasets, benchmarking it against the individual constituent models to demonstrate its enhanced predictive accuracy. Additionally, we conduct a sensitivity analysis to explore how varying perplexity weighting influences the ensemble’s performance, providing insights into optimal model selection strategies. To illustrate the practical utility of our approach, we present a case study using a proprietary viral dataset, highlighting ProPEC’s applicability in real-world scenarios. By addressing the current limitations of single PLM in predicting viral protein mutations, ProPEC serves as a robust and efficient tool that enhances our understanding of viral protein evolution. Ultimately, improving the accuracy of viral protein mutation predictions through optimal zero-shot model selection has the potential to accelerate antiviral drug discovery and strengthen our ability to respond to emerging viral threats.

## Methods

### Overview

ProPEC is an ensemble model designed for zero-shot viral protein mutant effect prediction, as outlined in Figure 1. It consists of a model selection module and a model integration module. The model selection module identifies the PLM with the lowest perplexity on the wild-type proteins, which is then used by the model integration module to score mutants. The final mutant fitness score is generated by combining this PLM’s predictions with those from the GEMME.

**Figure 1.**
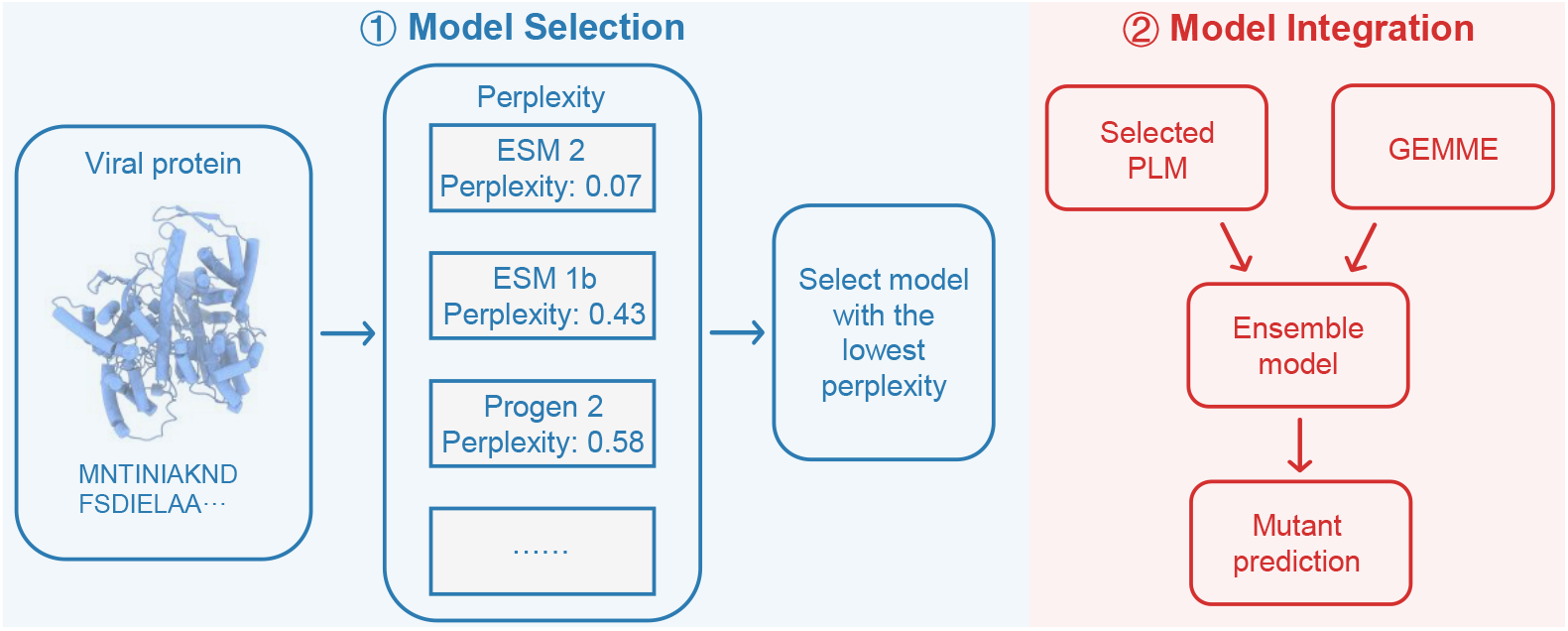
Overview of the ProPEC workflow. Initially, the wild-type sequence of the given viral protein is obtained. The perplexity of given sequence is then calculated using various deep learning models and subsequently normalized. The model with the lowest normalized perplexity is selected and integrated with GEMME. The ensemble model is then utilized to predict the fitness of the protein mutants.

### Model Selection Module

The model selection module select the most optimal PLM from a pool of candidates to serve as the primary predictor. We hypothesize that the lower the perplexity of a PLM has on a given protein, the more accurate its zero-shot performance is in predicting mutant fitness for that protein. This hypothesis is based on the definition of perplexity, which reflecting the model’s comprehension of the wild-type protein. A lower perplexity suggests a deeper understanding, which in turn leads to more accurate mutation effect predictions for given protein. (see more experimental support in **Results**)

Let *M* = {*M*_1_, *M*_2_, *M*_3_, …, *M*_*N*_ } represent the set of candidate PLMs, which includes models such as ESM2,^14^ ESM-1b,^29^ ProGen2,^15^ ESM-IF,^30^ and Tranception.^24^ We define *f* (*M, s*) →**R** as the perplexity of model *M* on protein sequence *s* (details on the calculation of perplexity for each model can be found in the Supporting Informations). For a given sequence *s*, the model with the lowest perplexity *M*_*k*_ is selected by :

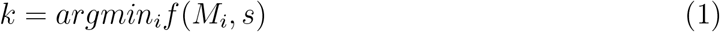

The selected *M*_*k*_ is then used as the input for the integration module to score the mutants.

### Model Integration Module

The model integration module scores all possible single-site mutants on the wild-type protein. First, each single-site mutant is scored using the selected PLM. Subsequently, the same mutants are scored by the MSA-based GEMME model. The final score is obtained by averaging the scores from both models.

### Evaluation Settings

We evaluated the performance of ProPEC on both a public dataset (ProteinGym) and an unpublished dataset (T7 RNA polymerase activity dataset). We applied the described workflow to score all possible single-site mutants for each protein. We then extracted those mutants with available experimental data and compared the predicted values to the experimental results using spearman correlation. A higher correlation indicates greater predictive accuracy of the model. Additionally, we evaluated the variance in spearman correlation across the datasets; a lower variance suggests greater robustness of the model.

### Datasets

We utilize a subset of ProteinGym,^24^ an extensive collection of deep mutational scanning (DMS) assays designed for the comparative evaluation of different models. Specifically, we focused on 23 viral protein datasets from the ProteinGym substitution benchmark. These datasets were selected based on the input length limitation of models such as ESM-1b, ^29^ which have a maximum input length of 1024 tokens. This selection process allowed us to retain a diverse yet manageable set of viral proteins for our analyses, while ensuring compatibility with the constraints of the computational models used. Additionally, to construct inputs for ESM-IF,^30^ we obtained the protein structures using AlphaFold2^31^ or downloaded them directly from AlphaFold DB when available. ^32^

T7 RNA polymerase activity dataset is dervied from the another manuscript currently under review.^33^ The dataset comprises 69 samples, including mutant sequences and their corresponding catalytic activities(Table S3). This dataset is used to validate ProPEC’s performance on viral protein mutations beyond those in ProteinGym. Permission for the use of this data was obtained from all relevant authors, in accordance with ethical guidelines.

### Metrics

To evaluate the predictive performance of ProPEC compared with other models, we utilized the spearman correlation coefficient (SPC) as our primary metric. It is defined as:

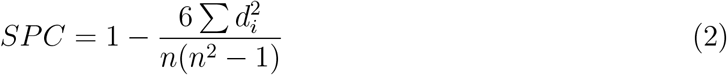

where *d*_*i*_ is the difference between the ranks of the predicted and experimental values for the *i*^*th*^ mutant, and *n* is the number of mutants. As higher SPC indicates that the model’s predictions are more closely aligned with the experimental rankings, reflecting greater accuracy. In addition to calculating the mean SPC across all datasets, we also computed the variance of SPC to assess the robustness of the model’s performance:

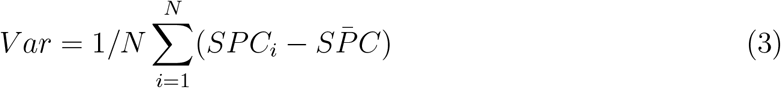

where *SPC*_*i*_ is the spearman correlation for the *i*^*th*^ dataset,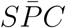 is the mean *SPC* acorss all datasets and *N* is the number of datasets. By combining both the mean SPC and its variance, we provide a comprehensive evaluation of the different models’ accuracy and reliability in predicting mutant fitness across diverse protein datasets.

## Results and discussion

To identify suitable models for inclusion in ProPEC, we systematically evaluate the relationship between the performance of various PLMs on 23 viral protein datasets from the ProteinGym benchmark and their perplexity on wild-type sequences. As shown in Figure 2, we observe that while some models do not exhibit outstanding predictive performance on mutation effects, their perplexity on viral wild-type proteins is strongly inversely correlated with their spearman correlation scores for mutant predictions. This inverse relationship forms the basis of our approach for selecting PLMs based on perplexity. Based on these two criteria: a pearson correlation between mutation effect predictions and wild-type perplexity less than –0.5 and a coefficient of determination (*r*^2^) greater than 0.2, we select five representative models as our candidate models for ensemble integration. These models, including ESM-IF,^30^ ESM-1b,^29^ ESM-2,^14^ Tranception,^24^ and ProGen2,^15^ are ranked and selected based on their perplexity on wild-type sequences of target proteins before being integrated with GEMME in the ProPEC ensemble (see Methods for details). The corresponding analysis for non-viral protein datasets has also been extended.(Figure 6) While a negative correlation between predictive performance and perplexity is still observed overall, the data distribution was more divergent, leading the correlation weaker compared to viral proteins.

**Figure 2.**
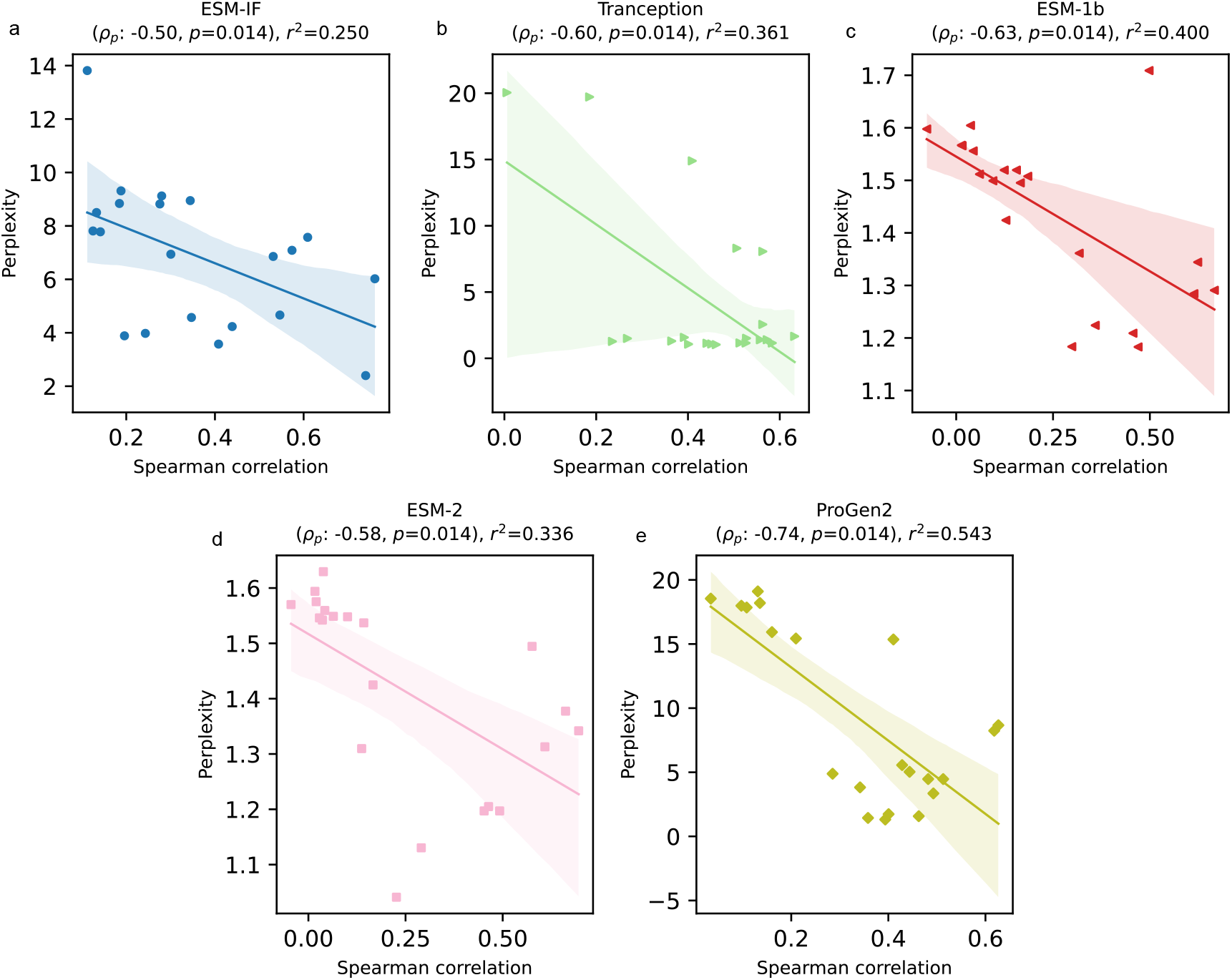
The performance of several PLMs on the viral protein dataset in ProteinGym shows a strong negative correlation between spearman scores and perplexity: (a) ESM-IF1, (b) Tranception, (c) ESM-1b, (d) ESM-2, and (e) ProGen2. These models have a Pearson correlation of less than -0.5 and an *r*^2^ greater than 0.2, making them suitable candidates for the integrated model.

We assess the performance of the ProPEC ensemble and its constituent models across 23 viral protein datasets (Figure 3a). On average, ProPEC significantly outperform all other baselines, including GEMME, demonstrating not only a higher average spearman correlation but also lower variance. In contrast, neither the naive averaging ensemble model nor the individual PLMs surpass GEMME in zero-shot performance. This outcome is not entirely unexpected, as PLMs may not consistently outperform MSA-based methods in predicting variant effects, despite their strong ability to model the statistical characteristics of protein sequences or structures.^24,34,35^ Notably, even when the integration step with GEMME is omitted, the ensemble of PLMs selected based on perplexity alone still outperform GEMME, coming in just behind the full ProPEC ensemble.This result underscores the effectiveness and stability of using perplexity as a criterion for model selection.

**Figure 3.**
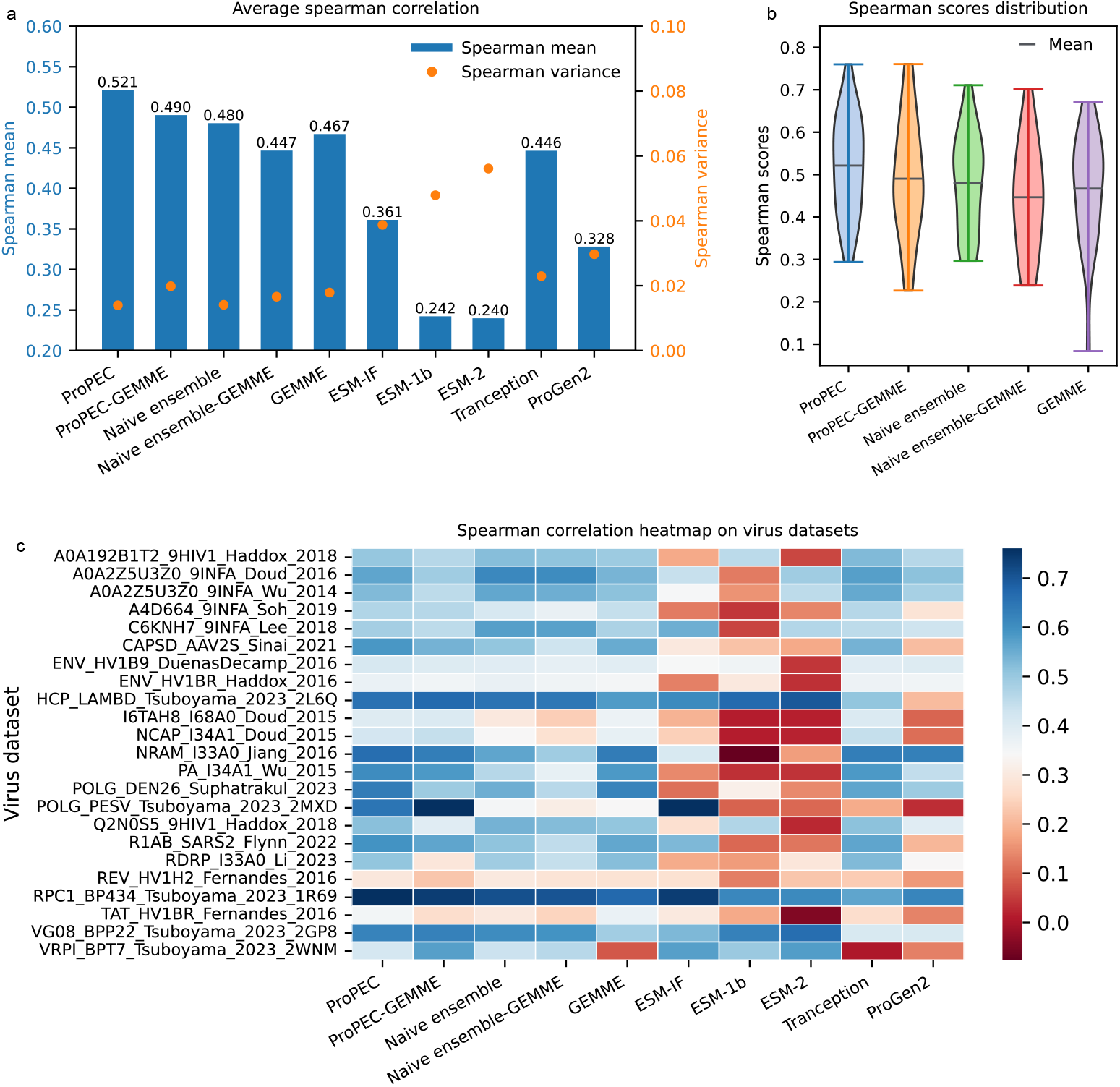
Overall performance on the ProteinGym viral protein mutation dataset. (a) The average performance of the integrated model and individual models was evaluated on 23 viral mutation datasets using the mean spearman correlation. ProPEC demonstrates significantly better performance. (b) Distribution of spearman correlation scores for the integrated models and GEMME across the 23 viral mutation datasets. (c) Heatmap showing the individual performance of the integrated models and GEMME on the 23 viral mutation datasets.

To further understand how ProPEC enhances the performance of the ensemble, we conduct additional statistical analyses on the models’ performance across the viral protein datasets. Compared to a naive averaging ensemble approach, the perplexity-based ensemble method not only elevates the upper limit of zero-shot performance but also mitigates the risk of lower-end outliers (Figure 3b,Table 2). Specifically, as shown in Figure 3c, no single model emerges as the best performer across all 23 datasets. However, when faced with individual datasets, the perplexity-based ensemble method effectively selects the most suitable models for ensemble, leveraging the strengths of different PLMs. Additionally, ProPEC’s integration of perplexity-selected PLMs with GEMME further enhances the applicability and robustness of the approach across diverse datasets.

**Table 1:** Perplexity summary of representive models.

| Metric | ESM-2 | ESM-1b | ProGen2 | Tranception | ESM-IF |
| --- | --- | --- | --- | --- | --- |
| Mean | 1.293 | 1.261 | 7.839 | 5.884 | 4.713 |
| Std | 0.127 | 0.153 | 5.388 | 5.887 | 2.207 |
| Median | 1.271 | 1.224 | 6.119 | 2.716 | 4.055 |
| Minimum | 1.080 | 1.025 | 1.438 | 1.057 | 1.887 |
| Maximum | 1.631 | 1.623 | 19.095 | 20.359 | 13.525 |

**Table 2:** Details of performance of the integrated models.

| Metric | ProPEC | ProPEC-GEMME | Naive ensemble | Naive ensemble-GEMME | GEMME |
| --- | --- | --- | --- | --- | --- |
| Mean | 0.521 | 0.490 | 0.480 | 0.447 | 0.467 |
| Variance | 0.014 | 0.020 | 0.014 | 0.017 | 0.018 |
| Maximum | 0.760 | 0.761 | 0.711 | 0.703 | 0.671 |
| Minimum | 0.294 | 0.227 | 0.297 | 0.239 | 0.084 |

Merging the benefits of model selection based on perplexity with the established performance of GEMME, ProPEC could serve as a powerful strategy for zero-shot prediction in complex protein variant effect prediction tasks.

To further refine our understanding of ProPEC’s performance, we conduct a parameter sensitivity analysis focusing on weight adjustments for base models with varying perplexity scores. Originally, ProPEC integrates only the base model with the lowest perplexity score into the ensemble. By introducing and varying a weight parameter *t*, we allow models with higher perplexities to contribute to the final prediction, with their influence controlled by the magnitude of *t*. The weight *w*_*i*_ of each candidate model *i* is computed as:

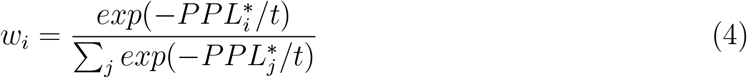

where 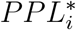 denotes the standardized perplexity of model *i*. As *t* increases, the differences in weights between models decreases, resulting in a more evenly distributed contribution from each model. Conversely, as *t* decreases, the ensemble’s reliance on the model with the lowest perplexity intensifies. This approach allowed us to perform a weighted averaging of the models’ predictions based on their adjusted perplexity scores.

Performance of ensemble models with different *t* is also evaluated by spearman correlation.(Figure 4a-c). Notably, before integrated with GEMME, introducing a small weight (*t* = 0.5) to the ensemble models with higher perplexity scores results in a modest improvement in the average spearman correlation, increasing it from 0.49 to 0.52. However, further increasing the perplexity weights to *t* = 1 does not lead to additional performance gains. By contrast, the average spearman correlation tends to decrease when the weight *t* increased to 2, indicating a potential limit to the benefits of incorporating less confident models. When these weighted models were further integrated with GEMME, their performance did not surpass the original ProPEC configuration (t=0), where only the model with the lowest perplexity was used. This suggests that the original configuration remains effective, even when considering the potential benefits of including additional models.Figure 4b illustrates the distribution of spearman correlation scores for different perplexity weight settings, both before and after integration with GEMME. Before integration, increasing the perplexity weight *t* generally made the ensemble’s performance more consistent and stable across datasets. This adjustment leads to higher minimum spearman scores and lower maximum scores, resulting in a higher mean value. This outcome is not surprising, as it reflects a trade-off similar to that observed in simple averaging ensemble methods: increasing the weight on additional models reduces the upper bound of performance but mitigates the risk of poor predictions. Considering the detailed breakdown of performance, including mean, variance, maximum, and minimum spearman correlations, the original ProPEC configuration (with *t* = 0) achieves a more effective balance between high accuracy and stability. Although moderate adjustments in perplexity weighting can yield slight improvements in performance, the benefits diminish as more weight is assigned to models with higher perplexity scores. This emphasizes the importance of careful model selection based on perplexity to ensure that the ensemble model remains both accurate and robust across diverse datasets.

**Figure 4.**
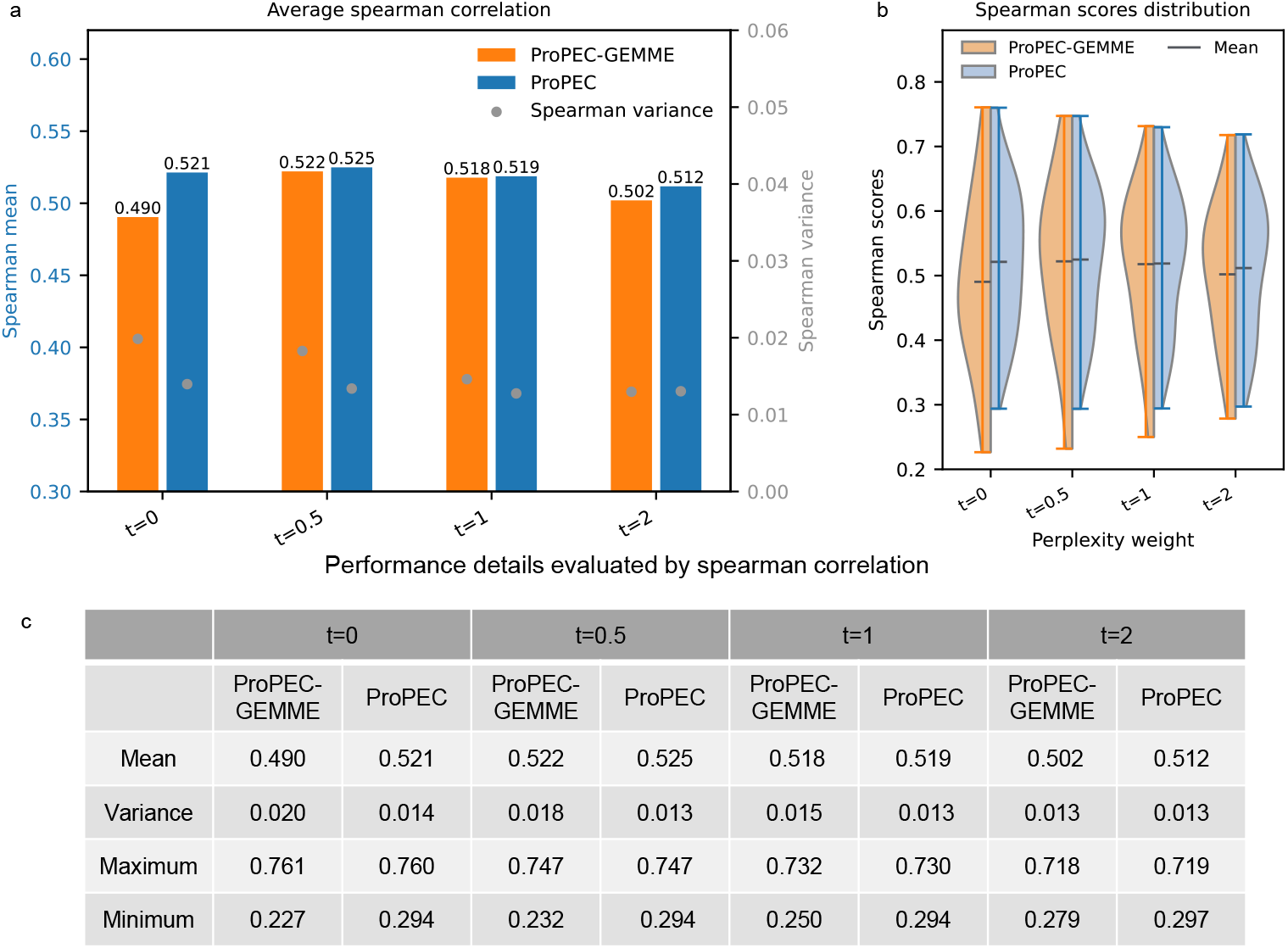
Comparison of integrated models under different perplexity weights. (a) The average spearman correlation was tested on 23 viral mutation datasets. (b) Distribution of spearman correlation scores across the 23 viral mutation datasets. (c)Table of performance details for models with different perplexity weights.

To further demonstrate the applicability and generalizability of ProPEC, we apply it to a proprietary T7 RNA polymerase activity dataset. T7 RNA polymerase, derived from bacteriophage T7, is a critical enzyme widely employed in in vitro transcription experiments^36,37^ and mRNA vaccine production,^38,39^ making its activity paramount to its industrial value.^40^ Consequently, enhancing the activity of T7 through directed evolution has been a focal point of research. Notably, T7 RNA polymerase is not included in the Proteingym benchmark datasets, presenting an ideal opportunity to test the robustness of ProPEC on previously uncharacterized proteins. In this case study, we utilize 69 single-site mutants to evaluate the performance of ProPEC. As shown in Figure 5a, the base models’ performance exhibits a negative correlation with their perplexity on the wild-type T7 sequence, consistent with our findings on Proteingym. Particularly, the ESM-IF model, which has the highest perplexity, displays a negative correlation between its predictions and the experimental measurements, indicating poor predictive performance. In stark contrast, the ESM2 model, which has the lowest perplexity, emerges as the top performer among the base models, underscoring the effectiveness of perplexity as a criterion for model selection in ProPEC.

**Figure 5.**
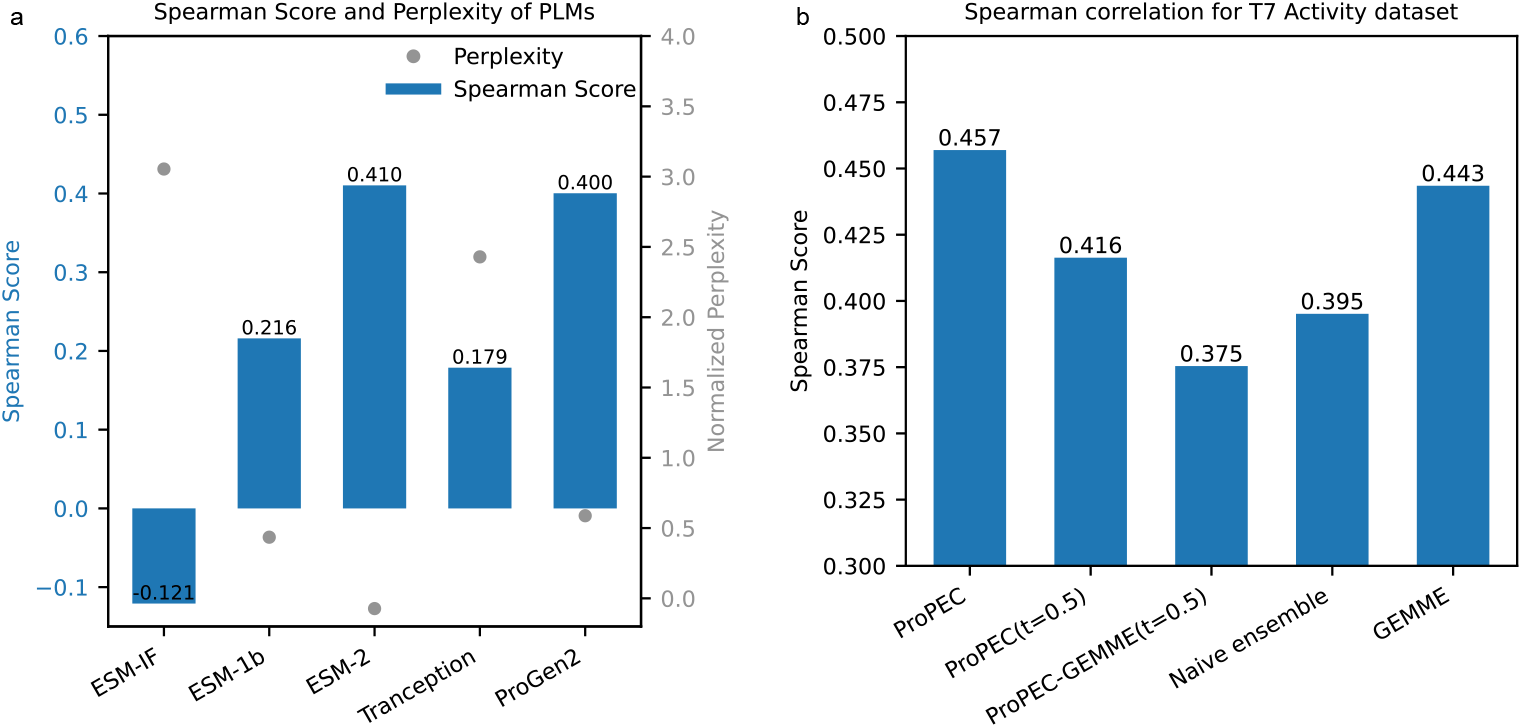
Figure 5: Performance on the T7 RNA polymerase activity dataset. (a) Comparison of the spearman correlation and the perplexity of the wild-type sequence for the PLMs. (b) Performance comparison between the integrated models and GEMME.

**Figure 6.**
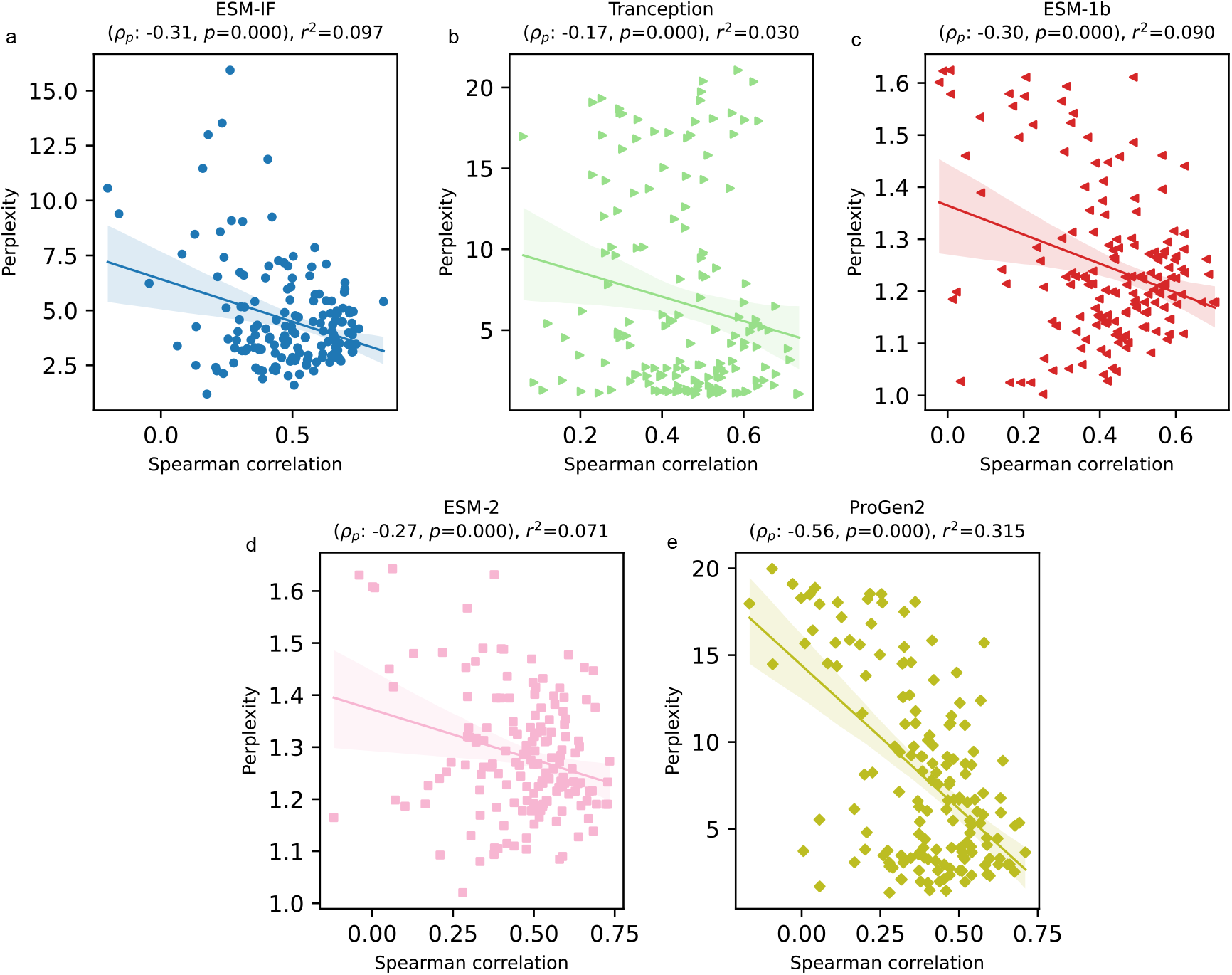
Figure S1: The performance of several PLMs on the non-viral protein dataset in ProteinGym : (a) ESM-IF1, (b) Tranception, (c) ESM-1b, (d) ESM-2, and (e) ProGen2.

Moving beyond individual models, we compare the performance of several ensemble models on the T7 dataset. Our ProPEC approach, which integrates perplexity-selected PLM with GEMME, achieved the highest spearman correlation score of 0.46. On the other hand, the naive averaging of all PLMs without considering perplexity (PLMs+GEMME) resultes in a spearman score of 0.40, highlighting the significance of model selection based on perplexity. Moreover,the ensemble’s performance declined when the moderate perplexity weight (t=0.5) applied, consistent with our parameter sensitivity analysis. These results demonstrate that ProPEC’s strategy of combining MSA-based information from GEMME alongside the most confident (i.e., lowest perplexity) PLMs strikes an optimal balance between accuracy and robustness.

## Conclusion

Predicting mutation effects in viral proteins is crucial for understanding viral evolution and developing antiviral strategies. In this work, we introduce ProPEC, an ensemble model that integrates multiple PLMs with the traditional GEMME approach, utilizing a perplexity-based model selection criterion. This approach enhances the accuracy and robustness of zero-shot predictions, particularly when dealing with the complex mutation landscapes characteristic of viral proteins.

Our extensive evaluations across 23 viral protein datasets demonstrated that ProPEC significantly outperforms individual PLMs and the GEMME model alone, achieving higher average spearman correlations and lower variance. These findings underscore the effectiveness of using perplexity as a criterion for model selection, enabling ProPEC to balance predictive accuracy and stability. By applying ProPEC to the T7 RNA polymerase activity dataset—an entirely unseen protein not included in the Proteingym benchmark—further we further validate the model’s generalizability and robustness.

The use of perplexity as a model selection strategy offers a new perspective on choosing among different models when faced with the challenge of predicting the mutant fitness with no prior data. Despite the our extensive evaluation of this approach, there are several opportunities for future work. While there is a general correlation between lower perplexity and better model performance, this relationship is not definitive. This suggests that while perplexity is a useful tool, it may not be the most decisive criterion for model selection. Future research could explore refining this approach. For example, developing more sophisticated selection methods that build upon perplexity and biological insights.

Based on the superior performance in predicting mutation fitness, ProPEC can effectively support the engineering and evolution of viral proteins. The integration of perplexity-based selection within ensemble modeling provides a more efficient preliminary screening tool and recommend single-site mutants for viral protein engineering and therapeutic design.

## Acknowledgement

This work was supported by the Computational Biology Key Program of Shanghai Science and Technology Commission (23JS1400600), Shanghai Jiao Tong University Scientific and Technological Innovation Funds (21×010200843), and Science and Technology Innovation Key R & D Program of Chongqing (CSTB2022TIAD-STX0017), the Student Innovation Center at Shanghai Jiao Tong University, and Shanghai Artificial Intelligence Laboratory.

## Supporting Information Available

The following file is available free of charge.

- Analysis on non-viral protein datasets.
- Detailed method for calculating perplexity.

## Analysis on non-viral protein datasets

In the analysis of non-viral protein datasets, the performance of five representative models also exhibits a negative correlation between prediction accuracy and perplexity.(Figure 6) This trend is consistent with findings in viral protein datasets. However, the absolute values of the pearson correlation coefficients and *r*^2^ values in non-viral datasets are generally lower. This indicates a weaker correlation.

## Detailed method for calculating perplexity

### ESM-2

One of the pre-training tasks of ESM-2 involves the recovery of original residues from substituted residues.^14^ ESM-2 can generate a probability distribution based on a wild-type sequence, reflecting the model’s expectations of what the sequence should be. Formally, a wild-type protein can be expressed as **R** = (*r*_1_, *r*_2_, …, *r*_*N*_), where each *r*_*i*_ ∈ *ℛ*^*V*^ represents the one-hot encoding of the *i*_*th*_ residue (1 for the true residue, 0 for all others), *V* denotes the residue vocabulary size, and *N* is the number of residues. The ESM-2 model produces a series of probability distributions 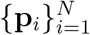 where 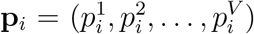 and 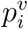 signifies the probability of residue *v* occurring at the *i*_*th*_ position. Then, the cross-entropy measures the difference between the predicted distribution and the actual residues. Specifically, the cross-entropy loss for a single residue prediction at the *i*_*th*_ position is calculated as:

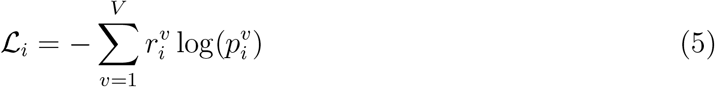

The total loss for the entire sequence **R** is computed as the sum of the individual losses over all positions:

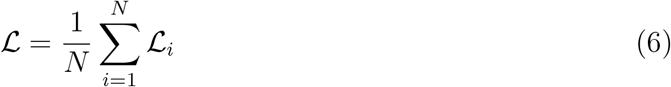

This loss estimates how well the model can predict the correct residue for each position in the sequence. The perplexity of the model, which indicates its uncertainty in predicting the correct residues, is then derived from the cross-entropy loss. Formally, perplexity (*PPL*) is calculated as:

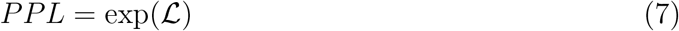

A lower perplexity value suggests that the model is more confident in its predictions, while a higher perplexity implies more uncertainty. In the context of ESM-2, the perplexity reflects the model’s ability to capture the underlying sequence patterns of the wild-type protein.

### ESM-1b

The perplexity calculation of ESM-1b ^29^ is the same as ESM-2.

### ESM-IF

ESM-IF is pre-trained with inverse folding, where the model generates a protein sequence based on a structure to which the sequence can be folded.^30^ We denote a protein structure as *S*. Based on *S*, the ESM-IF model auto-regressively generates a series of probability distributions 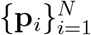where 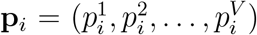 and 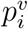 signifies the probability of residue *v* occurring at the *i*_*th*_ position. We also use cross-entropy to measure the difference between the predicted distribution and the actual residues:

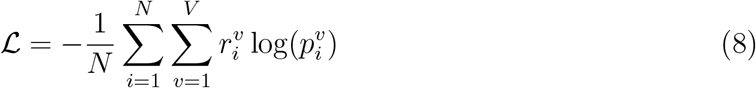

The perplexity of the model, which indicates its uncertainty in predicting the correct folded sequence, is then derived from the cross-entropy loss. Formally, perplexity (*PPL*) is calculated as:

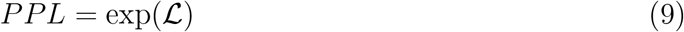

In the context of ESM-IF, the perplexity reflects the model’s ability to predict the correct sequence patterns of the wild-type structure.

### ProGen2

ProGen2 is a transformer-based auto-regressive model pre-trained to generate protein sequences.^15^ It generates a protein sequence **R** = (*r*_1_, *r*_2_, …, *r*_*N*_) by predicting the probability distribution for each residue *r*_*i*_ based on the previous residues (*r*_1_, *r*_2_, …, *r*_*i−*1_). Formally, the model generates a probability distribution **p**_*i*_ for each residue position *i*, where:

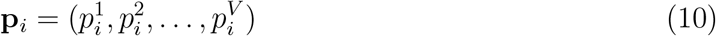

with 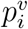 denoting the probability of residue *v* occurring at the *i*_*th*_ position in the sequence.

The sequence generation is auto-regressive, meaning that each predicted distribution **p**_*i*_ is conditioned on the previous predictions (*r*_1_, *r*_2_, …, *r*_*i−*1_):

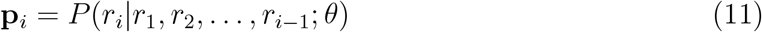

where *θ* represents the model parameters. The total loss across the sequence is computed as follows:

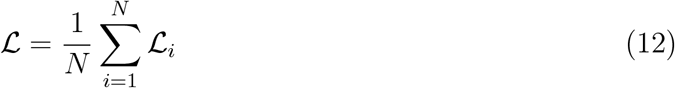

And the perplexity is derived from the cross-entropy loss:

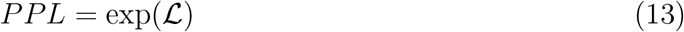

A lower perplexity score suggests that ProGen2 is more confident in its sequence generation, meaning it better understands the input sequence.

### Tranception

The perplexity calculation of Tranception^24^ is the same as ProGen2.

To facilitate a fair comparison of perplexity across different models, we standardize the perplexity values using the formula:

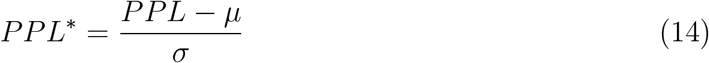

where *PPL*^∗^ represents the standardized perplexity and *PPL* is the raw perplexity score. *µ* is the mean and *σ* is the standard deviation of the perplexity scores across all the protein sequences on Proteingym. This standardization accounts for the inherent differences in perplexity calculation logic among the models, ensuring that subsequent comparisons are robust and meaningful.

